# *Ex vivo* midgut cultures of *Aedes aegypti* are efficiently infected by mosquito-borne alpha- and flaviviruses

**DOI:** 10.1101/2022.08.09.503233

**Authors:** Ana Lucia Rosales Rosas, Lanjiao Wang, Sara Goossens, Arno Cuvry, Li-Hsin Li, Nanci Santos-Ferreira, Alina Soto, Kai Dallmeier, Joana Rocha-Pereira, Leen Delang

## Abstract

*Aedes aegypti* mosquitoes can transmit several arboviruses, including chikungunya virus (CHIKV), dengue virus (DENV), and Zika virus (ZIKV). When blood-feeding on a virus-infected human, the mosquito ingests the virus into the midgut (stomach), where it replicates and must overcome the midgut barrier to disseminate to other organs and ultimately be transmitted via the saliva. Current tools to study mosquito-borne viruses (MBVs) include 2D-cell culture systems and *in vivo* mosquito infection models, which offer great advantages, yet have some limitations.

Here, we describe a long-term *ex vivo* culture of *Ae. aegypti* midguts. Cultured midguts were metabolically active for 7 days in a 96-well plate at 28°C and were permissive to ZIKV, DENV, Ross River virus (RRV) and CHIKV. *Ex vivo* midguts from *Culex pipiens* mosquitoes were found to be permissive to Usutu virus (USUV). Immunofluorescence staining confirmed viral protein synthesis in CHIKV-infected midguts of *Ae. aegypti*. Furthermore, fluorescence microscopy revealed replication and spread of a reporter DENV in specific regions of the midgut. In addition, two known antiviral molecules, β-D-N^4^-hydroxycytidine (NHC) and 7-deaza-2’-*C*-methyladenosine (7DMA), were able to inhibit CHIKV and ZIKV replication, respectively, in the *ex vivo* model.

Together, our results show that *ex vivo* midguts can be efficiently infected with mosquito-borne alpha- and flaviviruses and employed to evaluate antiviral drugs. Furthermore, the setup can be extended to other mosquito species. *Ex vivo* midgut cultures could thus be a new model to study MBVs, offering the advantage of reduced biosafety measures compared to infecting living mosquitoes.

**Importance:** Mosquito-borne viruses (MBVs) are a significant global health threat since they can cause severe diseases in humans, such as hemorrhagic fever, encephalitis, and chronic arthritis. MBVs rely on the mosquito vector to infect new hosts and perpetuate virus transmission. No therapeutics are currently available. The study of arbovirus infection in the mosquito vector can greatly contribute to elucidating strategies for controlling arbovirus transmission. This work investigated the infection of midguts from *Aedes aegypti* mosquitoes in an *ex vivo* platform. We found several MBVs capable of replicating in the midgut tissue, including viruses of major health importance, such as dengue, chikungunya, and Zika viruses. Additionally, antiviral compounds reduced arbovirus infection in the cultured midgut tissue. Overall, the midgut model emerges as a useful tool for diverse applications such as studying tissue-specific responses to virus infection and screening potential anti-arboviral molecules.

## 1. Introduction

Arboviruses are a diverse group of viruses that rely on arthropod vectors such as mosquitoes, ticks, and sandflies to infect susceptible vertebrate hosts and perpetuate transmission. Three arboviral families/orders are of most clinical relevance: *Bunyavirales, Flaviviridae*, and *Togaviridae*, as they encompass important arboviruses of global health concern, such as dengue virus (DENV), chikungunya virus (CHIKV), yellow fever virus, and Zika virus (ZIKV) (1). These four mosquito-borne viruses (MBVs) are mainly transmitted by female *Aedes* (*Ae*.) mosquitoes (e.g., *Ae. aegypti* and *Ae. albopictus*) (2), with *Ae. aegypti* mosquitoes considered highly competent vectors because of their anthropophilic behavior (3).

When blood-feeding on a virus-infected human or animal, a female mosquito will ingest the virus-containing blood in the midgut (i.e., mosquito stomach), where the virus must replicate and overcome the midgut barrier to subsequently disseminate to mosquito secondary organs. As the mosquito becomes systemically infected, resulting in a high viral load, the virus will reach the salivary glands. The virus will be released in the saliva upon the next bite, allowing virus transmission when the mosquito feeds on uninfected hosts (4, 5). The midgut is thus the initial site for infection and replication of any arbovirus in the mosquito. Virus replication in and spread across midgut cells are a requisite for a productive arbovirus infection in the mosquito (4).

*In vitro* cell cultures can be useful to study arbovirus infection and other relevant processes in mosquitoes; however, they present several limitations. Most of the established mosquito-derived cell lines are not derived from tissues relevant to specific stages of mosquito-virus interactions (e.g., salivary glands or midgut), but were generated from larval or embryonic tissue (e.g., *C6/36* [larvae] and *Aag-2* [embryos] cells) (6). Therefore, results obtained from *in vitro* experiments are ought to be taken cautiously. Furthermore, cell lines are considered homogenous cultures, both genetically and phenotypically, and single-cell type populations might not well represent multicellular tissues such as a midgut (7, 8). Moreover, some biological differences might arise among the same cell line coming from different laboratories (7).

Another method commonly used to study mosquito-virus interactions is the mosquito infection model. Through this approach, mosquito infection occurs artificially via a bloodmeal (membrane filled with infectious blood), and mosquitoes are sacrificed to assess viral infection and vector competence (9). Although living-infected mosquitoes provide valuable transmission data, the value of these data is directly proportional to the labor-intensive nature of the method. Transmission studies require skilled personnel for the handling of infected mosquitoes and facilities with a high biosafety level that ensure proper containment, and carry the risks of prick injury with virus-infected dissection tools or of infected mosquitoes escaping. Additionally, considerable variability among mosquito infection studies might exist as colonies bearing the same name can be highly genetically divergent among diverse laboratories (10).

*Ex vivo* organ cultures have been arising as an advantageous tool in research and have also been described for insects. For instance, *ex vivo* culture of insect organs has been successfully and widely defined for ixodid ticks. The synganglion, midgut, and salivary glands dissected from ticks were viable for 10 days and permissive to both Langat and Powassan viruses. (8, 11). On the contrary, reports on *ex vivo* organ cultures from mosquito tissues are scarce and mainly focused on germline and fat body tissues (12, 13). Despite the essential implication of the mosquito midgut in arbovirus replication and dissemination, no *ex vivo* mosquito midgut culture has been properly described. To date, only a short-term *ex vivo* assay in mosquito midguts has been reported, in which the midguts were only maintained for 35 hours post dissection in tissue culture plates (14).

Nevertheless, e*x vivo* cultures could offer a unique perspective to (i) study vector-virus interactions in a relevant tissue, for example, the mosquito midgut, (ii) reveal tissue-specific responses to virus infection, or (iii) study tissue-specific metabolic processes. Furthermore, *ex vivo* mosquito midgut cultures could provide a convenient and relevant platform to generate results that can be extrapolated to living mosquitoes. Additionally, *ex vivo* midgut cultures provide a controlled environment in which many parameters can be easily adjusted, and they constitute a safer option due to the low degree of containment required while working with BSL2/3 arboviruses, compared to working with live mosquito infection models.

On the lookout for novel tools that could be of use to study arbovirus infection within the arthropod vector, we dissected *Ae. aegypti* midguts and established their viability for a 7-day period. We also infected the midgut tissues with several mosquito-borne alpha- and flaviviruses, confirming the production of infectious viral particles. Furthermore, previously reported antiviral drugs were able to reduce arbovirus replication in the treated midguts. We thus developed an *ex vivo* mosquito midgut model that constitutes a valuable tool to study MBVs which complements the arsenal available to the mosquito arbovirus research field.

## 2. Results

### Viability of *Ae. aegypti* midguts cultured *ex vivo*

We first determined the viability of the dissected mosquito midguts in 96-well culture plates. Since motility could be observed along the gut tissue following the dissection, a bioassay was performed based on the number of contractions by the hindgut (Supplementary material, Movie 1). The same area of the hindgut was analyzed for all biological replicates (Fig. 1, b). Dissected midguts presented peristalsis in the hindgut for up to 7 days post dissection (d.p.d.) (Fig. 1, d), indicating that the tissue was viable. At 10 d.p.d., the mean frequency in the *ex vivo* cultured midguts was significantly reduced compared to day 0 measurements (day 0: 14.37 s vs day 10: 6.74 s) (Fig. 1, d), suggesting that the tissue’s viability had decreased.

**Figure 1.**
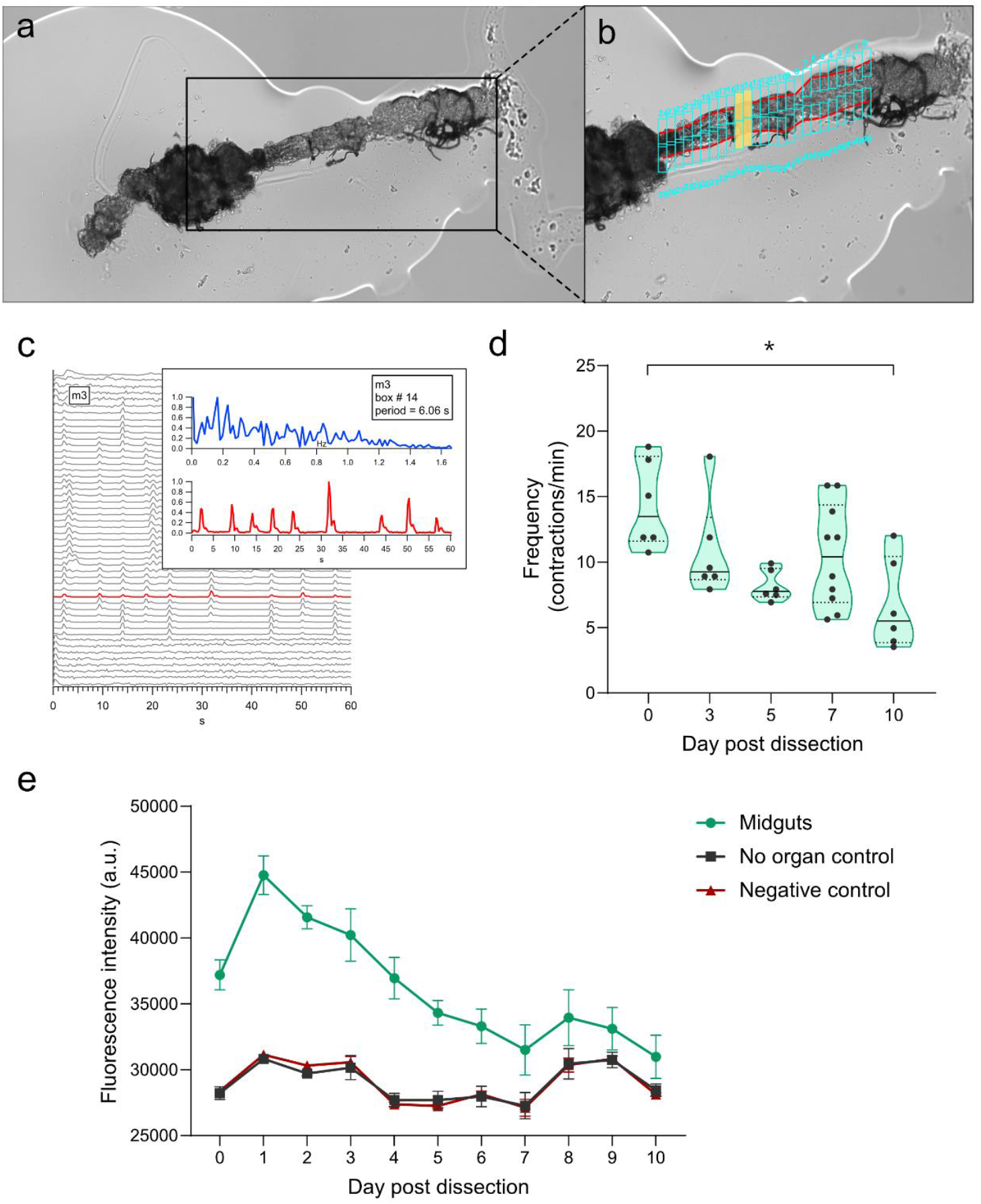
*Ex vivo* cultured mosquito guts were viable for 7 days post dissection, based on gut peristalsis and a resazurin-based assay. **a**, Representative image of a mosquito gut placed in a carboxymethyl cellulose (CMC) drop for videography. **b**, Close-up of the hindgut which comprises the area of analysis (AOA) manually set and outlined by the red lines. Cyan boxes denoted the regions to be analyzed along the AOA. The number and size of the regions (cyan boxes) were adjusted to be 25 boxes on each side of the hindgut. **c**, Representative kymograph showing the output for one mosquito gut analysis. Each line in the kymograph corresponded to an individual region, which was further investigated in a plot. In the figure, the red highlighted line in the kymograph correlates to the presented plot displaying a representative analysis readout for this region. **d**, Contractions observed per minute in function of time post dissection in days. Each point represents one individual mosquito midgut. The black line represents the median value. The graph shows data from three independent experiments. Statistical significance was assessed with a Kruskal-Wallis test. Significantly different values are indicated by asterisks: *, P <0.05. **e**, Viability readouts exhibit metabolic activity for a period of 10 days for tested midguts, no organ and negative controls. Assays were performed with 2 or 3 biological replicates, each replicate composed of 3 midguts per well, along with its corresponding controls. Error bars represent the standard errors of the means per time point. The graph shows data from three independent assays.

We further assessed the metabolic activity of the *ex vivo* cultured midguts in a resazurin salt-based assay. The dissected midguts remained metabolically active up until 7 d.p.d. (Fig. 1, e), corresponding with the data obtained from gut peristalsis. Fluorescence intensity in the tested midguts was still slightly higher than the “no organ” controls (NO, used to measure background signal) and negative controls (NC, paraformaldehyde fixated midguts) at 8, 9, and 10 d.p.d.. However, there was more variability among biological replicates and therefore 7 d.p.d. was selected as the end time point for following experiments.

Additionally, the effect of the neurotransmitter serotonin and the anticholinergic drug atropine on peristalsis was assessed to rule out that this visual feature might be a reflex. Midguts (day 0 post dissection) were exposed to either serotonin, atropine, or midgut medium (mock-exposure), after which their gut peristalsis was recorded. The mean frequency displayed by the mock-exposed midguts was 7.349 s (5 biological replicates, median value: 7.48 s). Midguts exposed for 1 hour to serotonin displayed a slight increase in gut motility, with a mean peristaltic period of 12.76 s (6 biological replicates, median value: 10.61 s), whereas atropine-exposed midguts exhibit a somewhat reduced peristalsis, with a mean peristaltic period of 6.72 s (6 biological replicates, median value: 7.58 s). Despite the observed trends, there were no statistically significant differences among the conditions evaluated (Supplementary Fig. S1).

### Infection of dissected *Ae. aegypti* midguts with mosquito-borne alpha- and flaviviruses

Next, *ex vivo* virus infection was performed in the cultured midguts to test their capacity to support the replication of several arboviruses. Replication kinetics were evaluated for Ross River virus (RRV) and CHIKV as representative members of the alphavirus genus of the *Togaviridae* family. Both viruses were able to efficiently replicate in the *ex vivo* cultured midguts with an increase in viral RNA levels between 1 and 3 days post infection (d.p.i.), compared to viral RNA levels at 2 hours p.i. (h.p.i.) (Fig. 2, a). Infectious virus titers of RRV and CHIKV increased accordingly in the midgut tissue (Fig 2, b), while no infectious virus was observed yet at 2 h.p.i. (data not shown). The peak of viral RNA and viral titer levels was detected at 3 d.p.i. for both alphaviruses.

**Figure 2.**
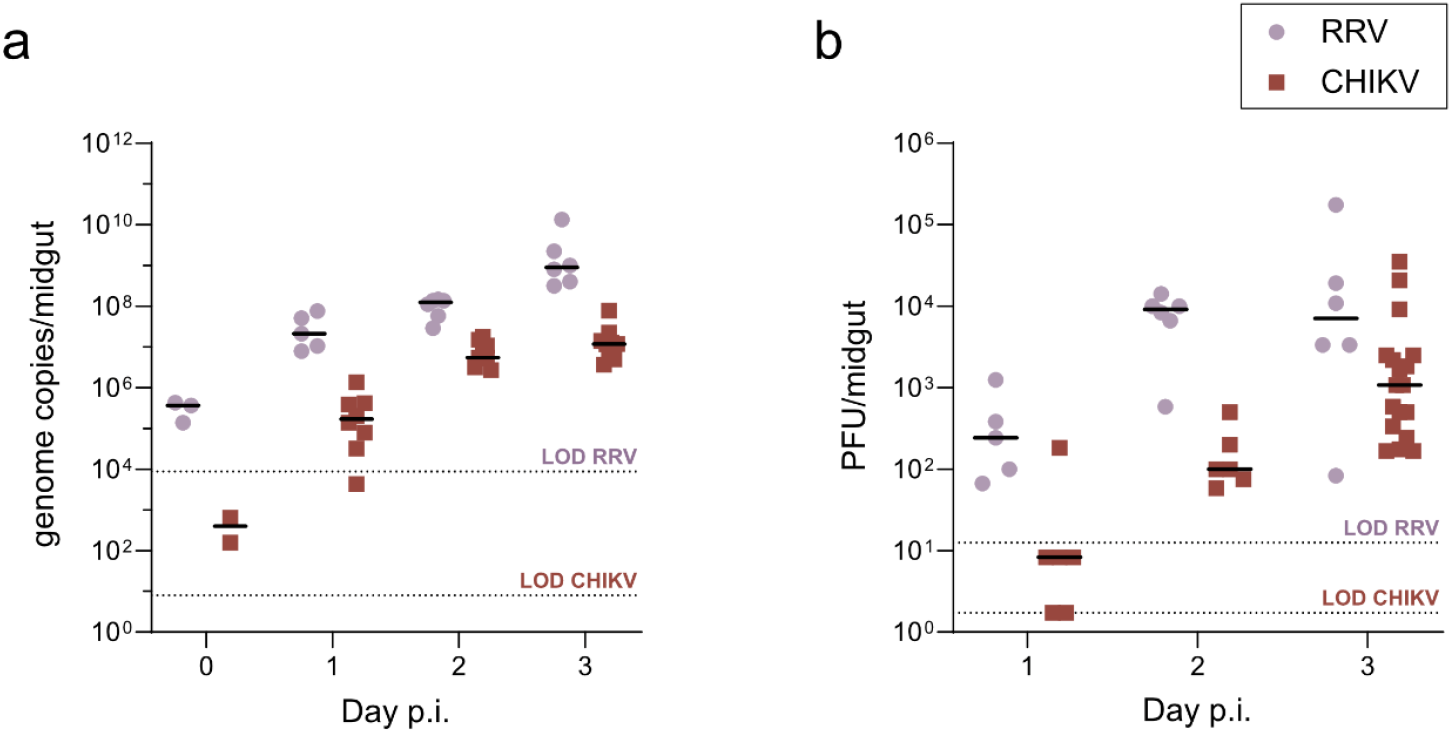
*Ex vivo* cultured midguts supported Ross River (RRV) and chikungunya virus (CHIKV) infection. **a**, Viral RNA levels in the mosquito midguts were quantified at 2 hours (day 0), 1, 2 and 3 days p.i. by means of qRT-PCR. **b**, Infectious virus loads in the mosquito midguts were quantified by means of plaque assay. Each dot represents an individual midgut organ. The black line represents the median value. Data correspond to at least two independent replication kinetics assays. LOD: Limit of detection of the corresponding assay.

To evaluate whether *Aedes*-borne flaviviruses could replicate in the *ex vivo* midguts, the replication kinetics of DENV-2 and ZIKV were studied. As both viruses needed a longer period to replicate in the midgut tissue (data not shown), viral RNA and infectious virus levels were measured starting at 5 d.p.i.. Viral titers were increased at 5 d.p.i. and further peaked at 7 d.p.i., with DENV-2 reaching RNA levels up to 2.5 × 10^8^ genome copies/midgut and a viral titer of 21 PFU/midgut. On the contrary, ZIKV replication and infection presented more variability among individual midguts. The maximum amount of ZIKV RNA and infectious virus quantified at 7 d.p.i. were 2.1 × 10^6^ genome copies/midgut and 4.2 × 10^2^ PFU/midgut, respectively (Fig 3).

**Figure 3.**
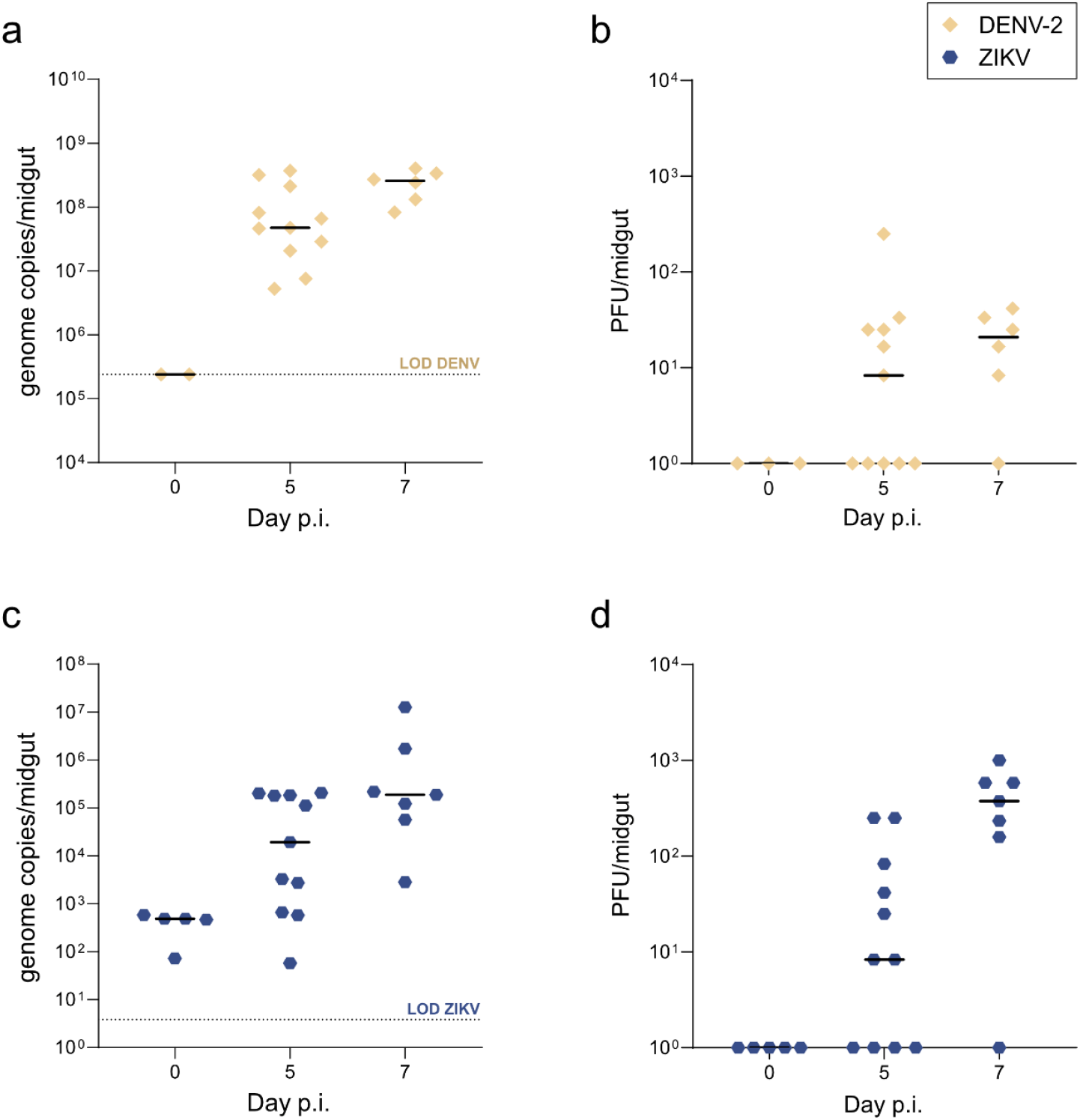
*Ex vivo* cultured midguts supported dengue virus serotype 2 (DENV-2) and Zika virus (ZIKV) infection. **a, c**, Viral RNA levels in the mosquito midguts were quantified at 2 hours, 5 and 7 days p.i. by means of qRT-PCR. **b, d**, Infectious virus loads in the mosquito midguts were quantified by means of plaque assay. Each symbol represents an individual midgut organ. The black line represents the median value. Data correspond to two independent replication kinetics assays. LOD: Limit of detection of the corresponding assay.

### Usutu virus (USUV) can modestly replicate in *ex vivo* cultured midguts from *Culex pipiens* mosquitoes

As a proof-of-concept, the *ex vivo* midgut model was applied to another mosquito species: *Culex (Cx*.*) pipiens*. The midguts of *Cx. pipiens* thrived in culture and showed peristaltic movements during their incubation, as *Ae. aegypti* midguts did. Therefore, we evaluated the susceptibility of these *Cx*. midguts to USUV, as *Cx. pipiens* mosquitoes are considered a competent vector for this virus (15). USUV infection was assessed in the midguts at 7 d.p.i. by qRT-PCR and plaque assay. Only 50% (3 out of 6) of the midguts became infected with USUV, reaching up to 1.8 × 10^6^ genome copies/midgut. However, only 2 out of these 3 USUV-infected midguts contained infectious virus, amounting to 6.4 × 10^2^ PFU/mL. Of note, a considerable amount of viral RNA was detected in samples corresponding to 2 h.p.i. and in 2 out of the 3 negative controls included in the assay, yet no infectious virus was detected in these samples (Fig. 4, a, b).

**Figure 4.**
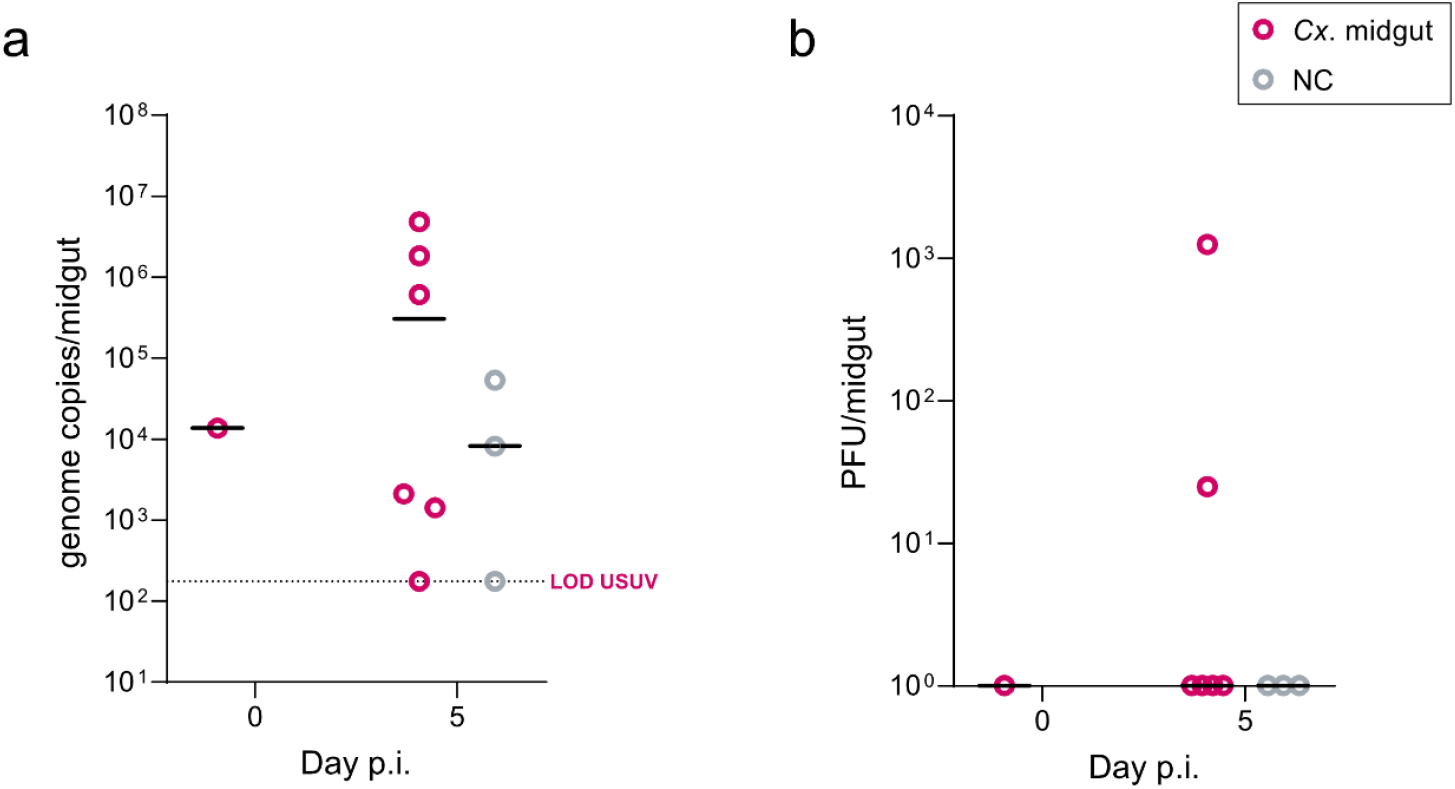
*Ex vivo* cultured midguts were partially permissive to Usutu virus (USUV) infection. **a**, Viral RNA levels in the mosquito midguts were quantified at 2 hours and 5 days p.i. by means of qRT-PCR. **b**, Infectious virus loads in the mosquito midguts were quantified by means of plaque assay. Each symbol represents an individual midgut organ. The black line represents the median value. LOD: Limit of detection.

### CHIKV viral protein synthesis in infected *ex vivo* midguts

Previous infection experiments indicated the susceptibility of the *ex vivo* midguts to infection with several arboviruses. To further corroborate these results, the CHIKV E2 glycoprotein was visualized in CHIKV-infected midguts at day 3 p.i. by immunostaining. Specific staining for this viral protein could be observed in the infected guts, while no signal was observed in the negative controls (fixated guts that followed the same infection protocol; Fig. 5, a). At day 3 p.i., E2 protein synthesis was mainly detected and spread along the posterior midgut region (Fig. 5, b, c), with some infection foci located in the hindgut region (Fig. 5, c). The E2 signal was also present in the tracheal tubes that remained attached to the midgut after dissection (Fig. 5, b, arrow heads).

**Figure 5.**
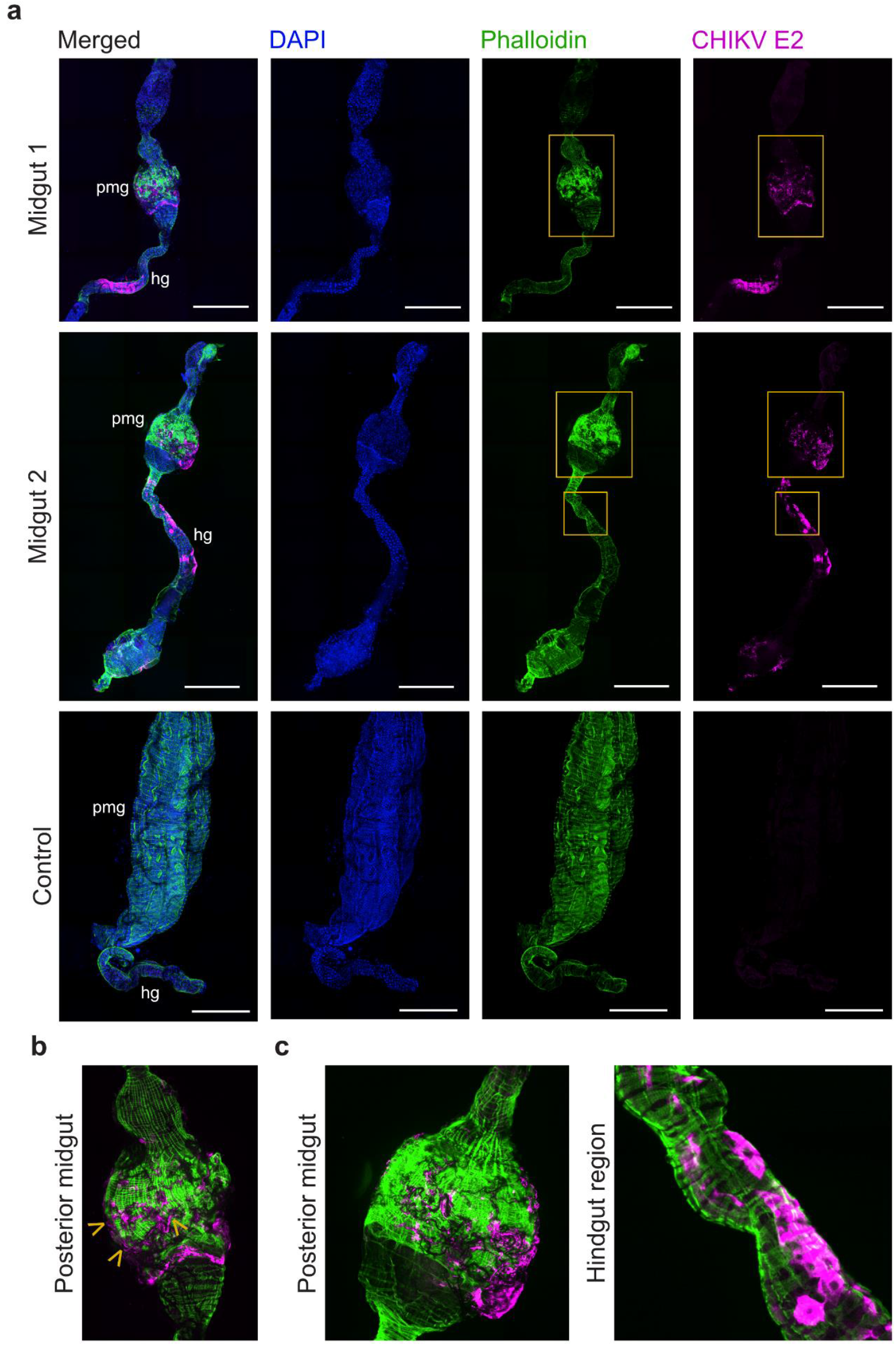
Detection of CHIKV protein synthesis in infected *ex vivo* midgut cultures. **a**, Magnification, 25X. Overlay of blue, green, and magenta filter imaging are shown for two infected midgut organs and one negative control midgut. Imaging of each individual channel as follows: in blue, DAPI-stained cell nuclei; in green, actin filaments stained with phalloidin; and in magenta, E2 viral protein. No E2 expression was detected in the negative control midguts. The scale bar in panels, represented by the white line, corresponds to 400 µM. pmg: posterior midgut. hg: hindgut. **b**, Close-up panel showing infection foci in the posterior midgut region corresponding to the yellow squares indicated in **a** for “Midgut 1”. Arrow heads indicate E2 expression localized in tracheal tubes of the midgut. **c**, Close-up panel showing infection foci in the posterior midgut and hindgut region corresponding to the yellow squares indicated in **a** for “Midgut 2”.

### Infection of *ex vivo* cultured midguts with an mCherry-expressing DENV-2

To follow the progression of virus infection in the midguts by imaging, cultured mosquito midguts were infected with DENV-2 expressing the red fluorescent protein mCherry (DV2/mCherry,(16)) and imaged using the FLoid™ Cell Imaging Station (Life Technologies) at selected time points p.i.. The mCherry signal was used as a proxy for infection. DENV-2 mCherry infection was observed initially at day 3 p.i. as single or few foci in the posterior midgut region (Supplementary Fig. S2, a). Over time, these focal infection points increased in number and spread to neighboring areas, primarily along the posterior region of the midgut and the hindgut (Supplementary Fig. S2, b-e). No mCherry expression was observed in the negative control or mock-infected midguts at day 7 p.i. (Fig. 6, a). Following 7 days of infection, the mCherry signal had spread mostly along the posterior midgut region, forming infection foci comprising multiple cells (Fig. 6, b, c). Moreover, some infected cells were found in the tracheal tubes that remained in the midgut tissue after dissection (Supplementary Fig. S2, h).

**Figure 6.**
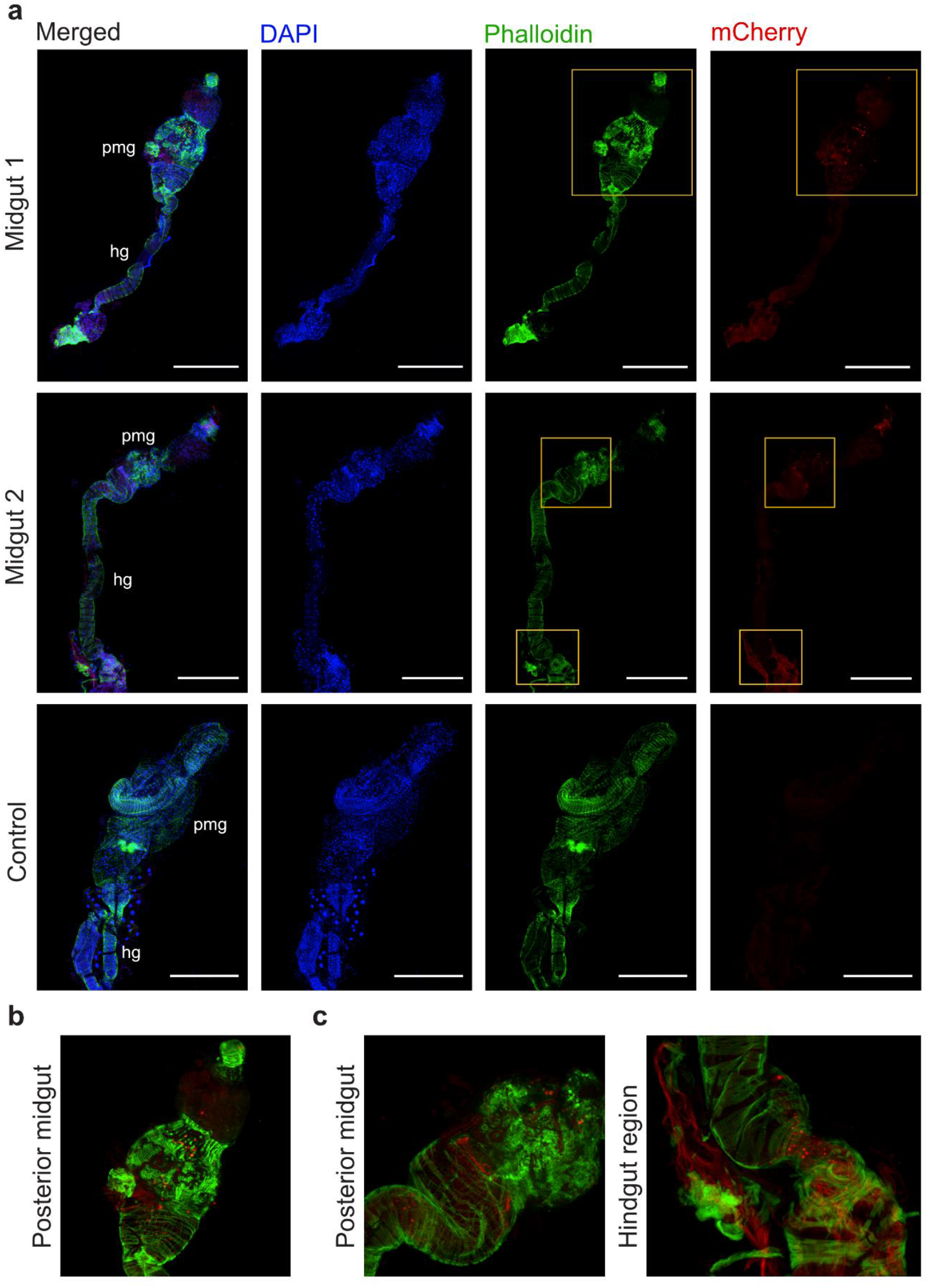
Replication of DENV-2 expressing mCherry in the *ex vivo* cultured midguts. **a**, Magnification, 25X. Confocal microscopy displays the DENV-2 infection in the *ex vivo* midguts at day 7 p.i., as seen by the mCherry (red) signal. Overlay of blue, green, and red filter imaging are shown for two midgut organs and one negative control midgut. Imaging of each individual channel as follows: in blue, DAPI-stained cell nuclei; in green, actin filaments stained with phalloidin; and in red, mCherry signal. No mCherry expression was detected in fixated or mock-infected midguts. The scale bar in panels, represented by the white line, corresponds to 400 µM. pmg: posterior midgut. hg: hindgut. **b**, Close-up panel showing several infection foci in the posterior midgut region corresponding to the yellow squares indicated in **a** for “Midgut 1”. **c**, Close-up panel showing infection foci in the posterior midgut and hindgut region corresponding to the yellow squares indicated in **a** for “Midgut 2”.

### Arbovirus replication is reduced upon treatment with antiviral drugs in midgut cultures

We next assessed whether the *ex vivo* mosquito midguts could be used to evaluate the antiviral activity of inhibitors against arboviruses. To this end, the antiviral activity of β-D-N^4^-hydroxycytidine, also known as EIDD-1931 or NHC, was tested against CHIKV at a concentration of 50 µM. At day 3 p.i., no difference in the CHIKV RNA levels was observed between the untreated and NHC-treated groups (Fig. 7, a). In contrast, NHC significantly reduced the infectious virus levels by 1 log (mean values, control: 1.0 × 10^4^; 50 µM NHC: 1.1 × 10^3^ PFU/midgut) (Fig. 7, b).

**Figure 7.**
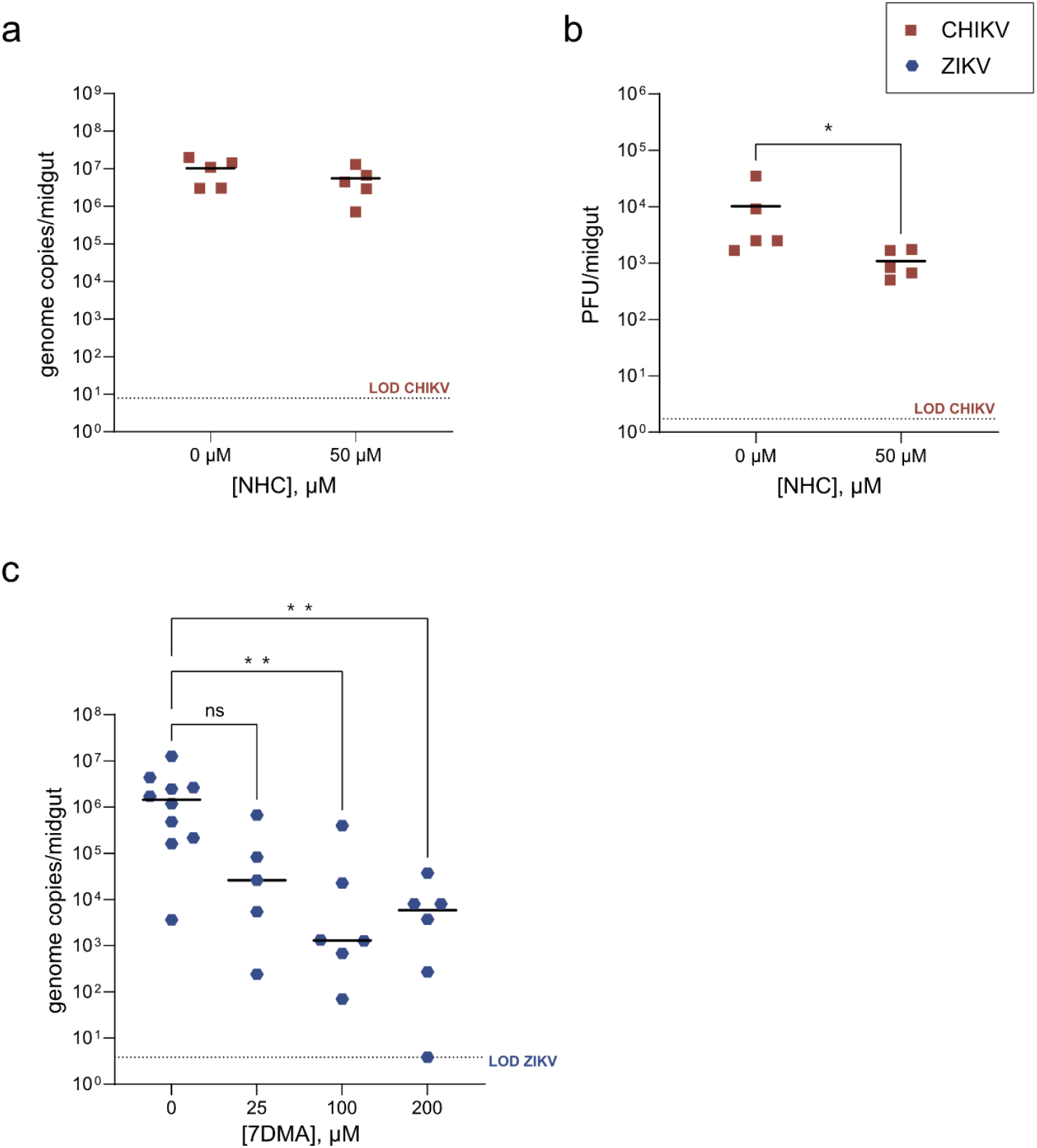
Antiviral activity of NHC and 7DMA in the *ex vivo* cultured mosquito midguts. **a**, CHIKV RNA loads in the mosquito midguts were quantified at day 3 p.i. by qRT-PCR. **b**, CHIKV infectious virus loads were quantified at day 3 p.i. by plaque assay. Statistical significance was assessed with a Mann-Whitney test. Significantly different values are indicated by asterisks: *, P<0.05. **c**, ZIKV RNA levels in the mosquito midguts were quantified at day 7 p.i. by qRT-PCR. Statistical significance was assessed with a Kruskal-Wallis test. Significantly different values are indicated by asterisks: **, P<0.05. ns: not significant. Each dot represents an individual midgut. The black line represents the mean value. LOD: Limit of detection of the corresponding assay.

As the *ex vivo* cultured midguts also supported flavivirus infection, the viral polymerase inhibitor 7-Deaza-2’-*C*-Methyladenosine (7-DMA) was tested against ZIKV. Viral RNA levels were reduced for all treated midguts (25, 100, and 200 µM of 7-DMA) compared to the untreated group at day 7.p.i.. However, only the groups treated with 7-DMA at 100 and 200 µM showed a significant difference compared to the untreated group, with 1.5 log and 2.4 log reductions (virus control: 2.6 × 10^6^; 100 µM: 7.0 × 10^4^; 200 µM: 9.5 × 10^3^ mean genome copies/midgut), respectively (Fig. 7, c).

## 3. Discussion

The study of arbovirus infection in the mosquito vector relies mainly on a combination of *in vitro* and *in vivo* approaches, which has yielded great progress in the knowledge regarding arbovirus biology, virus-vector interactions, and vector competence. While convenient and handy, *in vitro* cell culture systems have the shortcoming of not being as biologically relevant as mosquito infection models, aside from the limited selection of mosquito cell lines available. On the other hand, working with living infected mosquitoes grants useful information when studying virus-vector interactions or vector competence for a specific virus, but such *in vivo* models are not always accessible and require the implementation of cumbersome safety measures ensuring adequate containment while working with BSL2/3 pathogens (17). *Ex vivo* organ culture methods have not yet been described in-depth for the study of MBVs. Hence, in this study, we have established an *ex vivo* mosquito midgut model using dissected midguts from *Ae. aegypti* mosquitoes. When cultured *ex vivo*, the midguts displayed peristaltic contractions, mainly observed as waves along the hindgut, which is a frequent occurrence for this type of tissue (18). Here, we used this visual feature as a proxy for viability, paired with a custom script for gut motility in zebrafish (19) to measure the peristaltic period (time between contractions) in the *ex vivo* cultured guts, which remained stable for 7 days p.d.. This observation was further corroborated when measuring the metabolic activity of the dissected midguts over time in culture with PrestoBlue™ (PB). This resazurin-based reagent has been successfully used to assess cell viability and cytotoxicity in both two-dimensional cell monolayers and 3D cellular interfaces (including organ explants) (20–23). Here, we found that only after 7 hours of incubation of the *ex vivo* midguts (using multiple midgut organs per well) in presence of the PB reagent yielded a colorimetric change.

The *ex vivo* cultured mosquito midguts supported infection with four arboviruses, despite the unusual route of infection (through incubation with virus inoculum) that differs from what occurs naturally in mosquitoes (infection via blood feeding). Both viral RNA and infectious virus particles were detected in the mosquito midgut organs following a progressive increase over time, which also confirmed the viability of the midguts. These replication kinetics results indicated the preservation of midgut tissue in an *ex vivo* culture set-up and, more specifically, the presence of midgut cells that could constitute the target for arbovirus infection.

With *Ae. aegypti* being the main vector for both DENV-2 and ZIKV, it was however unexpected that infectious virus titers for both flaviviruses detected in the *ex vivo* midguts did not reach higher levels than what was inoculated (1 × 10^4^ PFU/mL), in contrast to viral RNA levels. Kinetics of DENV replication in mosquitoes have reported a steady increase in infectious virus until 8 d.p.i., after which it normally declined without affecting viral RNA levels (24, 25). Such results could explain the discrepancy seen between flavivirus RNA and infectious levels. More importantly, it has been described that within 2 hours of infection with DENV or ZIKV (both *in vivo* and *ex vivo*), there is a rapid induction of apoptosis in the *Ae. aegypti* midgut epithelium in an attempt of the host to control flavivirus infection (26). Nonetheless, this process might cause tissue damage, which correlates with the high midgut cell turnover rate in mosquitoes during a bloodmeal digestion (27). As such, we cannot disregard that the *ex vivo* midguts in our experiments undergo the same process during DENV or ZIKV infection, consequently damaging the tissue and thus limiting flaviviral replication. This hypothesis may also apply to *ex vivo* midguts of *Cx. pipiens* mosquitoes, as a comparable replication pattern can be observed when assessing USUV (flavivirus) infection at day 7 p.i.. Further addition of an apoptosis inhibitor to the virus inoculum used to infect the *ex vivo* midguts might elucidate whether rapid induction of apoptosis in the midgut epithelium is indeed a limiting factor for flavivirus replication in this setup.

Active virus replication was confirmed for CHIKV through an immunofluorescence assay. Envelope (E2) protein synthesis was localized in the posterior midgut region, where it would also occur when the mosquito ingests a CHIKV-infectious bloodmeal (28), regardless of the infection method employed with the *ex vivo* cultured midguts. Moreover, CHIKV infection of tracheal tubes in the tissue could be observed, consistent with *in vivo* reports (28). Similar results were obtained when analyzing the progression of DENV2/mCherry infection in the *ex vivo* midguts. Previously, DENV-2 infection in *Ae. aegypti* (Chetumal strain) midguts was reported to start with individual infected epithelial cells detected as early as 2 d.p.i., slowly progressing to infection foci consisting of multiple cells until the whole midgut organ was infected at 7-10 d.p.i. (24). In agreement with this, a less strong but akin infection pattern was observed in DENV2/mCherry-infected midguts, with few foci of several infected cells by 7 d.p.i.. Together, these data show that arbovirus infection occurred at considerable levels in *ex vivo* cultured midguts and therefore it could be used to study other aspects of arbovirus infection and facilitate the collection of preliminary data before experimenting with mosquitoes *in vivo*.

The use of antiviral compounds to inhibit virus infection in the mosquito vector is an innovative concept that might reduce or block arbovirus transmission from mosquitoes to humans. This idea involves antiviral molecules being ingested by adult mosquitoes when they take a bloodmeal on a mammalian host undergoing antiviral treatment. As the mosquito midgut is the entry point and key replication site for MBVs, studying the effect of antiviral compounds in the midgut tissue would be of great interest. For this purpose, the antiviral activity of two inhibitors was assessed in the *ex vivo* mosquito midguts. NHC is a nucleoside analog that has been characterized as a potent antiviral drug against alphaviruses *in vitro*, including CHIKV and Venezuelan equine encephalitis virus (VEEV) (29, 30). Various assays point to the compound acting as a pyrimidine analog that may target the viral polymerase domain of nsP4, provoking chain-termination (31). In addition, NHC also induces a high level of mutations in virus-specific RNAs, resulting in lethal mutagenesis (32). Consistent with these findings, CHIKV RNA levels were not reduced in NHC-treated midguts, but there was a marked decrease in virus infectivity. The viral polymerase inhibitor 7DMA has shown potent anti-ZIKV activity *in vitro* and delayed disease progression in mice (33). In the *ex vivo* mosquito midguts, 7DMA significantly reduced ZIKV RNA loads. Although the RNA levels presented some variability among the ZIKV-infected midguts, the inhibitory effect of 7DMA in the *ex vivo* mosquito midguts was significant and followed a dose-response relationship. These results indicate that *ex vivo* midgut cultures could be used to rationally select potential arbovirus-blocking molecules to be tested in living mosquitoes at a later stage.

In summary, we have established a long-term *ex vivo* mosquito midgut culture. To support this model, we have (a) assessed the viability of the *ex vivo* cultured midguts, (b) determined the replication kinetics of two alphaviruses (RRV and CHIKV) and two flaviviruses (DENV-2 and ZIKV), (c) detected viral protein synthesis and followed live arbovirus infection in infected midguts, (d) demonstrated that the *ex vivo* protocol can be translated to other mosquito genera, and, lastly, (e) evaluated the antiviral activity of two inhibitors against CHIKV and ZIKV. Altogether, we have provided a reference for the use of *ex vivo* mosquito midguts as a tool to study arbovirus infection and related processes, while offering groundwork for developing other *ex vivo* mosquito organ cultures that can potentially provide insightful data, like salivary glands.

## 4. Materials and Methods

### 4.1. Cells

#### Mammalian cells

African green monkey kidney cells (Vero cells, ATCC CCL-81) and Vero E6 cells (ATCC CRL-1586) were cultured in minimum essential medium (MEM 1X) enriched with 10% Fetal Bovine Serum (FBS), 1% sodium bicarbonate, 1% L-glutamine and 1% non-essential amino acids (NEAA). Baby hamster kidney cells (BHK, ATCC CCL-10) were maintained in Dulbecco’s Modified Eagle’s Medium (DMEM) containing 10% FBS, 1% sodium bicarbonate and 1% L-glutamine. Mammalian cell cultures were incubated at 37°C, with 5% CO_2_.

#### Mosquito cells

*Ae. albopictus* larval cells (C6/36, obtained from ATCC, CRL-1660) were maintained in Leibovitz’s L-15 medium containing 10% FBS, 1% Penicillin-Streptomycin (PenStrep), 1% NEAA, and 1% HEPES buffer. Mosquito-derived cell lines were incubated at 28°C, without CO_2_.

For cell culture assays containing virus or virus-infected material, the concentration of FBS in the medium was reduced to 2%, for both mammalian and mosquito cells. All cell culture media and supplements were obtained from Gibco™, ThermoFisher Scientific (Aalst, Belgium).

### 4.2. Viruses

#### Flaviviruses

Dengue virus serotype 2 (DENV-2/TH/1974, isolated in 1974 from human serum collected in Bangkok, Thailand, GenBank MK268692.1) was kindly provided by Prof. A. Failloux (Institut Pasteur, Paris, France) (34); ZIKV (SL1602, Suriname strain, GenBank KY348640.1) was acquired via the EVAg consortium (https://www.european-virus-archive.com). The infectious clone DENV-2 pDVWS601 used for the construction of the DENV-2 reporter virus expressing the red fluorescent protein mCherry (DV2/mCherry, New Guinea C strain, NGC, GenBank AF038403.1) was kindly provided by Prof. Andrew Davidson (University of Bristol, Bristol, UK) (16).

#### Alphaviruses

Ross River virus (RRV) was received from the National Collection of Pathogenic Viruses (UK; catalog number 0005281v); and CHIKV (Indian Ocean strain 899, GenBank FJ959103.1) was generously provided by Prof. Drosten (University of Bonn, Bonn, Germany) (35).

Virus stocks were prepared by passaging the isolates on Vero (for CHIKV and USUV) or C6/36 cells (for ZIKV, DENV-2, and RRV). Viral titers of the stocks were determined via plaque assay or end-point titration on Vero or BHK cells.

### 4.3. Compounds

7-Deaza-2’-*C*-Methyladenosine (7-DMA) was purchased from Carbosynth (Berkshire, UK) and dissolved in DMSO. β-D-N^4^-hydroxycytidine (EIDD-1931 or NHC) was purchased from MedChemExpress (Monmouth Junction, NJ, USA) and dissolved in DMSO.

### 4.4. *Ae. aegypti* rearing

*Ae. aegypti* Paea (Papeete, Tahiti, collected in 1994) were obtained via the Infavec2 consortium. For each rearing, eggs were hatched in dechlorinated tap water. After hatching, groups of ±400 larvae were transferred into trays containing 3 L of dechlorinated tap water and fed every day with a yeast tablet (Gayelord Hauser, Saint-Genis-Laval, France) until the pupae stage. Pupae were placed in small plastic containers inside cardboard cups for their emergence. Adult mosquitoes were supplied with cotton balls soaked in a 10% sucrose solution supplemented with 100 U/mL and 100 µg/mL of PenStrep. Cardboard cups containing adults were maintained at 28 ± 1 °C with a light/dark cycle of 16/8 h and 80% relative humidity.

### 4.5. *Cx. pipiens* rearing

*Cx. pipiens* biotype *pipiens* were kindly provided by Prof. Sander Koenraadt (Wageningen University & Research, Wageningen, Netherlands) (15). For each rearing, eggs rafts were hatched in trays containing 2 L of Milli-Q water (Synergy® UV, Merck, Germany). Larvae were fed continuously until the pupae stage with TetraMin® baby fish food (Tetra, Spectrum Brands, Germany). Pupae were collected as described in 4.4. Cardboard cups containing adults were maintained at 25 ± 1 °C with a light/dark cycle of 16/8 h and 70% relative humidity.

### 4.6. Mosquito midgut dissection and *ex vivo* culture

Unfed, antibiotic-treated female mosquitoes (3 – 7 days old) were cold-anaesthetized and surface sterilized by soaking in 70% ethanol for 20 s followed by soaking in Dulbecco’s phosphate-buffered saline (PBS) for 20 s. Next, mosquitoes were dissected in PBS on a petri dish using the stereomicroscope (VisiScope®, VWR). In brief, the midgut was exposed by carefully pulling the second to last segment of the mosquito abdomen. The gut of the mosquito was excised in its entirety (foregut, anterior and posterior midgut, and hindgut) to keep the tissue of interest (midgut) from degradation, and tracheal tubes were removed as much as possible without damaging the tissue. Following dissection, mosquito guts were washed twice in midgut medium, consisting of: Leibovitz’s L-15 medium supplemented with 2% FBS, 100 U/mL of PenStrep, 50 µg/mL of kanamycin, and 0.25 µg/mL of amphotericin B (Sigma Aldrich, USA), and finally placed in a 96-well tissue culture plate with a clear bottom (PerkinElmer®, USA) containing midgut medium. One mosquito gut was placed in each testing well. Plates were maintained at 28°C, without CO_2_. Of note, handling and optimization of the *ex vivo* mosquito midguts culture are extensively described in Supplementary Appendix 1.

### 4.7. Videography of gut peristalsis and analysis

*Ex vivo* cultured mosquito midguts were recorded with the Leica DMi8 microscope to quantify the peristalsis observed *ex vivo*. A drop of carboxymethyl cellulose (CMC) 0.8% diluted in Leibovitz’s L-15 medium was deposited on a microscope glass slide and one mosquito midgut was soaked in the drop. Forceps were used to gently arrange the hindgut in a position suitable for analysis (horizontally positioned on the slide). Each hindgut was recorded for a total of 1 minute 7 s (1 frame/0.3 s) and further discarded. This procedure was repeated at several time points during incubation starting from day 0 (right after dissection) to assess the contractibility, and hence viability, of the *ex vivo* cultured midguts over time.

For the analysis of the gut peristalsis, frames generated for each midgut were processed in IgorPro (Wavemetrics, USA) using a custom script described for the measurement of zebrafish gut motility (19). This script quantifies the changes in pixel intensity in a designated area of analysis as it sequentially goes through all frames generated during the video recording. As output, the software indicates individual peristaltic periodicity (seconds in between contractions) per point of evaluation and overall averages of peristaltic periodicity for each midgut analyzed. These outputs were annotated manually to check for artifacts that could be generated by debris or tracheal tubes.

Serotonin hydrochloride (Sigma Aldrich, USA) and atropine sulfate salt monohydrate (Sigma Aldrich, USA) were used to test the responsiveness of the mosquito midgut tissue. Midguts were incubated with either serotonin (20 µM), atropine (20 µM), or midgut medium for 1 hour, after which they were video recorded for further analysis.

### 4.8. PrestoBlue™ (PB) assay

The metabolic activity of the dissected midguts was further assessed using a resazurin salt-based cell viability reagent PrestoBlue™ (Invitrogen, USA). For these experiments, three dissected midguts were placed per well in a 96-well tissue culture plate. Each trio of midguts was considered a biological replicate. Using more than one midgut per well was necessary to ensure both a robust readout and a relatively rapid change of color in the testing medium. Midguts fixated in 4% paraformaldehyde (PFA, Sigma Aldrich, USA) were included in the assay as negative controls (NC). No organ (NO) controls were composed of wells where one tip of the forceps was dipped in during dissection.

The live midguts and the corresponding controls were tested with PB every day for a period of 10 days. In brief, the medium was carefully removed from the wells and replaced with 100 µL of a 1:10 dilution of PB prepared in midgut medium (without phenol red). Next, the midguts and their controls were incubated at 28°C without CO_2_ for 7 hours. Once the incubation time was completed, the PB-containing medium was transferred to a 96-well tissue culture plate and fluorescence intensity was measured at wavelengths 560 nm excitation and 590 nm emission with a Spark® Multimode Microplate Reader (Tecan Trading AG, Switzerland). New midgut medium was added to wells containing midguts and they were returned to the incubator (28°C, no CO_2_) until the next time point assessment. Two independent assays were carried out with at least 3 biological replicates per time point.

### 4.9. Infection and antiviral assays in *ex vivo* mosquito midguts

Mosquito guts were dissected as described in 4.6 and placed into a 96-well tissue culture plate filled with midgut medium. After removing the medium, a total of 1 × 10^4^ PFU/mL of the virus was added to each well. Of note, inocula of 1 × 10^5^ PFU/mL and 1 × 10^6^ PFU/mL were used for DENV-2 and USUV, respectively. Negative controls consisted of dissected guts fixated in 4% PFA for 30 minutes. Midguts were incubated for two hours at 28°C, without CO_2_. The virus inoculum was carefully removed, and midguts were washed two times with midgut medium. Fresh medium was finally added to all wells for incubation at 28°C, without CO_2_.

To test the activity of antiviral drugs in the *ex vivo* mosquito midguts, compound dilutions were prepared in midgut medium and added to the wells containing midguts to be treated, after which they were infected with 1 × 10^4-5^ PFU/mL of virus. Following two hours of incubation at 28°C, without CO_2_, the midguts were washed twice before adding midgut medium alone (for virus control midguts) or containing compound (for treated midguts). Midguts were returned to the incubator at 28°C, without CO_2_.

Incubation time for alphavirus-infected midguts was 3 days, while flavivirus-infected midguts were maintained for 7 days. Midguts were collected at several time points after infection for further analysis.

### 4.10. Determination of viral RNA levels and detection of infectious virus replication

Collected midguts were homogenized individually in 250 µL of PBS using bead disruption (2.8 mm beads, Precellys). The midgut homogenate was filtered using 0.8 µm MINI column filters (Sartorius, Germany) to remove debris, bacteria, and fungi. The filtered homogenate was used for further viral RNA isolation and qRT-PCR to determine viral RNA levels, and plaque assay to assess infectious virus particles.

Viral RNA isolation was performed with the NucleoSpin RNA Virus kit (Macherey-Nagel, Germany) following the manufacturer’s protocol. The sequences of primers and probes used for each virus are compiled in Table 1. One-Step, quantitative RT-PCR was performed for CHIKV and ZIKV in a total volume of 25 µL, consisting of 13.94 µL of RNase free water (Promega, USA), 6.25 µL of master mix (Eurogentec, Belgium), 0.375 µL of each forward and reverse primer (to a final concentration for each primer: 150 nM [CHIKV and USUV]; 900 nM [ZIKV]), 1 µL of probe (to a final concentration of 400 nM [CHIKV and USUV]; 200 nM [ZIKV]), 0.0625 µL of reverse transcriptase (Eurogentec, Belgium), and 3 µL of RNA sample. For RRV, the reaction mixture was prepared in a total volume of 20 µL, containing 5.2 µL of RNase free water, 10 µL of SYBR Green master mix (BioRad, USA), 1 µL of each forward and reverse primer (to a final concentration of 125 nM for each primer), 0.3 µL of reverse transcriptase (BioRad, USA), and 4 µL of RNA sample. For DENV-2, the reaction mixture was prepared to a final volume of 20 µL, consisting of 3 µL of RNase free water, 10 µL of SYBR Green master mix (BioRad, USA), 0.3 µL of each forward and reverse primer (to a final concentration of 900 nM for each primer), 0.4 µL of reverse transcriptase (BioRad, USA) and 6 µL of RNA sample.

**Table 1.**
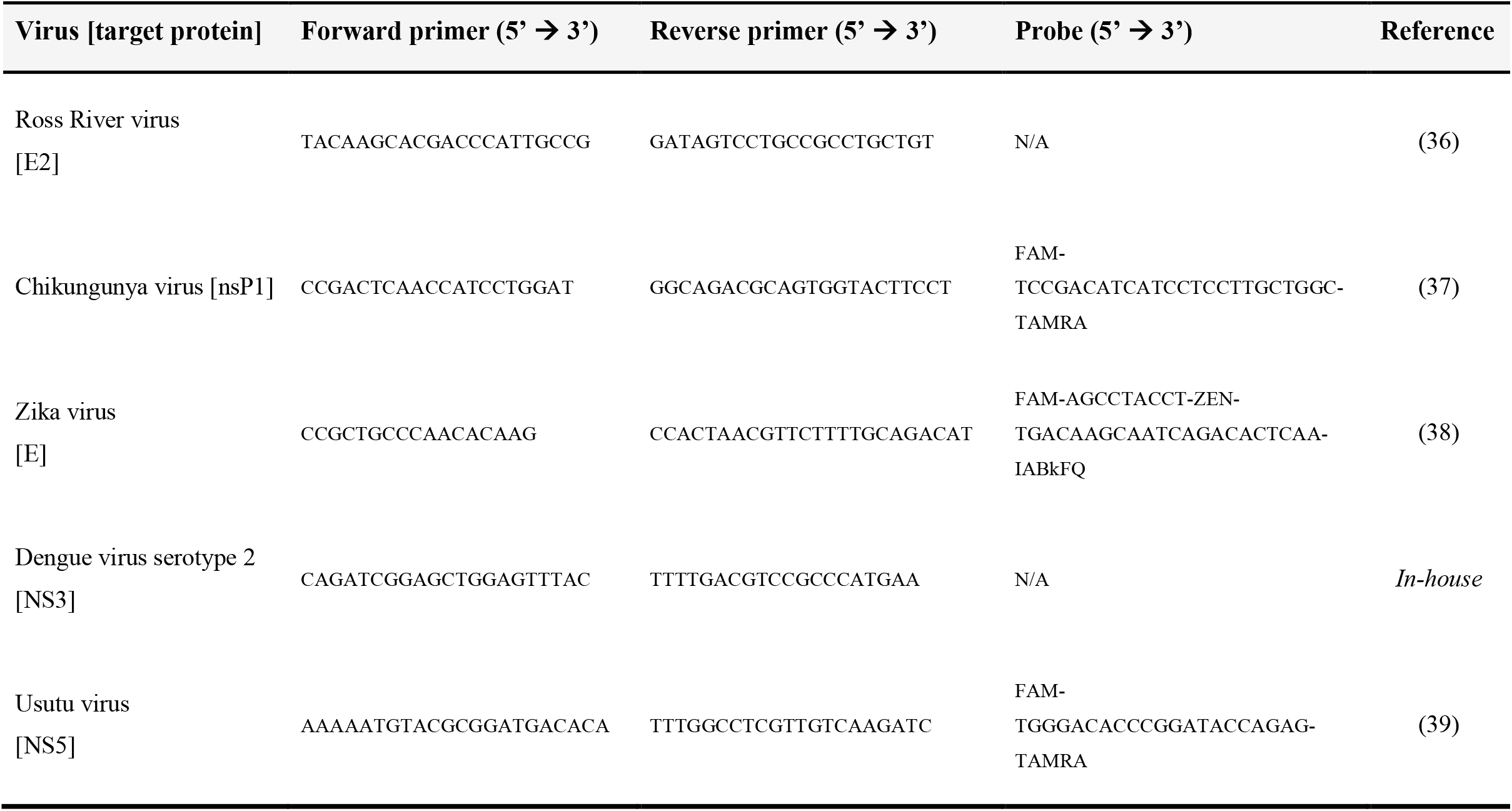
Sequences of primers and probes for qRT-PCR used in this study.

The qRT-PCR assays were performed using the QuantStudio™ 5 Real-Time PCR System (ThermoFisher Scientific, USA) with the following cycling conditions for CHIKV, ZIKV, and USUV: 30 min at 48°C, 10 min at 95°C, followed by 40 cycles of 15 s at 95°C, and 1 min at 60°C (55 °C for USUV). Cycling conditions for RRV were as follows: 10 min at 50°C, 3 min at 95°C, followed by 40 cycles of 15 s at 95°C, and 30 s at 60°C. For DENV-2, the following cycling conditions were used: 30 min at 48°C, 3 min at 95°C, followed by 40 cycles of 15 s at 95°C, and 1 min at 62°C.

For absolute quantification, standard curves were generated each run using 10-fold serial dilutions of: cDNA of CHIKV-nsP1, viral RNA isolated from RRV and DENV-2 Bangkok virus stocks or synthesized gBlocks™ gene fragments (Integrated DNA Technologies, USA) for ZIKV and USUV.

Plaque assays were performed to quantify infectious virus particles from infected midgut samples. Mammalian cells (Vero cells for RRV and CHIKV; BHK cells for ZIKV, DENV-2, and USUV) were seeded at a density of 2.5×10^5^ cells per well in 24-well tissue culture plates. One-day confluent monolayers of cells were inoculated with serial 10-fold dilutions of midgut samples, dilutions were prepared in 2% assay medium, and the inoculum was allowed to infect the cells for 2 hours at 37°C, with 5% CO_2_. A negative control consisting of 2% assay medium used for preparing the dilutions was included in all plaque assays.

The inoculum was removed and replaced with CMC 0.8% overlay diluted in RPMI medium (supplemented with 1% HEPES, 1% sodium bicarbonate, 1% L-glutamine, 1% PenStrep and 2% FBS). Infected cells were incubated for 3 (*Alphaviruses*) or 7 (*Flaviviruses*) days. After incubation, 1 mL of 4% PFA was added to each well on top of the overlay medium and allowed for fixation for 1 hour, after which the overlay mixture was discarded, and the wells were carefully washed with water. The culture plates were allowed to air dry, and then the wells were stained with 1% crystal violet solution (Sigma Aldrich, USA). Plaques were counted and titer of each sample was calculated.

### 4.11. Immunofluorescence staining of CHIKV viral proteins and imaging

Mosquito guts were dissected and infected with CHIKV as described in 4.6 and 4.9, respectively. At day 3 p.i., midguts were fixated with 4% PFA for one hour at room temperature. Fixated midguts were transferred to a µ-slide 18-well (Ibidi GmbH, Germany) plate and rinsed with PBS five times, for 5 minutes each time. Midguts were then saturated with PBS-T (0.1% Triton X-100 [Sigma Aldrich, USA] and 1% bovine serum albumin [BSA, Sigma Aldrich, USA] in PBS 1X) for 2 hours at room temperature. Primary antibody anti-E2 protein [Chk265] (Absolute Antibody, Wilton, UK) was diluted in PBS-T (1:500) and added to the fixed midguts for overnight incubation at 4°C. After incubation, midguts were rinsed five times with PBS-T, for 5 to 10 minutes each rinse, followed by incubation with Alexa Fluor 594 donkey anti-rabbit secondary antibody (ThermoFisher Scientific, USA, A-21207, diluted in PBS-T, 1:500) for 2 hours at room temperature. Midguts were rinsed five times with PBS-T, for 5 to 10 minutes each rinse. Phalloidin (diluted in 1% BSA in PBS 1X solution, final concentration 100 nm) incubation followed for 10 minutes, after which DAPI (diluted in PBS 1X, final concentration 100 nm) was added on top of the phalloidin solution and allowed to incubate for 10 minutes. Finally, the phalloidin/DAPI solution was removed, and midguts were rinsed with PBS. Midgut samples were mounted in microscope slides using Glycergel® mounting medium (Agilent, USA) and allowed to dry before imaging with a Leica DMi8 microscope (Leica Microsystems, Germany) and the Andor Dragonfly Confocal Spinning Disk microscope (Oxford Instruments, UK). All incubation steps were carried out on rotation, at 300 rpm.

### 4.12. Infection of *ex vivo* cultured midguts with DENV-2 expressing mCherry and imaging

Mosquito guts were dissected and infected with a DENV-2 expressing mCherry (16) as described in 4.6 and 4.9, respectively, using an inoculum of 1 × 10^5^ PFU/mL. Negative controls and mock-infected midguts were included. Mock-infected midguts were incubated in midgut medium instead of virus dilution. Midguts were monitored every day during the incubation period and checked under the FLoid™ Cell Imaging Station (Life Technologies) for any signal that indicated mCherry expression. At day 7 p.i., the mosquito midguts were fixated with 4% PFA for one hour at room temperature. After fixation, midguts followed the protocol described in 4.11, skipping the addition of any primary and secondary antibodies.

## 5. Acknowledgements

Special thanks to Prof. Johan Neyts (University of Leuven, Belgium) for allowing us to use his laboratory space, equipment, and compounds for our experimental work. We are grateful to Prof. Julia Dallman and Prof. Mason Klein (University of Miami, USA) for kindly providing the custom IgorPro script used for gut motility analysis. Many thanks to Prof. Sarah Merkling and Elodie Couderc (Institut Pasteur, Paris, France) for their helpful advice regarding the immunostaining of mosquito midguts. We would also like to thank the European virus archive, EVAg, supported by the European Union’s Horizon 2020 research and innovation program under grant agreement No. 871029, for supplying the ZIKV SL1602, Suriname strain. Thanks to Prof. Anna Bella Failloux (Institut Pasteur, Paris, France) for providing the DENV-2/ Bangkok strain. We thank Prof. Christian Drosten (University of Bonn, Germany) for providing the CHIKV 899 strain. We also thank Prof. Andrew Bristol (University of Bristol, UK) for providing the infectious clone DENV-2 pDVWS601, which served to construct the reporter DV2/mCherry used in this study. Many thanks to Prof. Pedro Elias Marques (University of Leuven, Belgium) for granting us the use of the Andor Dragonfly Confocal Spinning Disk microscope. Our sincere thanks to Prof. Sander Koenraadt (Wageningen University & Research, Netherlands) for kindly sharing *Cx. pipiens* mosquito egg rafts. Finally, we thank the Infravec2 Consortium for providing the *Ae. aegypti* mosquito eggs. This project was funded by KU Leuven (C22/18/007 and starting grant STG/19/008).

## Supplementary material

**Appendix I**. Handling and optimization of *ex vivo* mosquito midguts.

**Figure S1**. *Ex vivo* cultured mosquito guts were not significantly affected by external incubation with serotonin or atropine.

**Figure S2**. Replication of DENV-2 expressing mCherry in the *ex vivo* cultured midguts. **Figure S3**. ZIKV RNA loads in the mosquito midguts when cultured in medium or when using a foam substrate for support.

**Figure S4**. CHIKV replication kinetics in mosquito midguts cultured in medium and in carboxymethyl cellulose (CMC).

**Movie 1**. Representative mosquito hindgut videography used for the gut motility analysis. Peristalsis is observed as waves along the hindgut region.

