## Supplementary Appendix 1 for "*Ex vivo* midgut cultures of *Aedes aegypti* are efficiently infected by mosquito-borne alpha- and flaviviruses"

### Appendix 1. Handling and optimization of *ex vivo* mosquito midguts

Thorough optimization has been carried out to make the *ex vivo* mosquito midgut model an accessible and convenient tool for the study of arboviruses in the mosquito vector. Initially, mosquito midguts were placed in the well of a 96-well tissue culture plate, in which they remained suspended in culture medium. However, certain procedures (washing steps) were challenging due to the small size of the midguts and the high probability of pipetting out the tissue. Therefore, other support systems were tested to immobilize the midguts and facilitate their handling to overcome this limitation.

To this end, Matrigel® (Corning®) domes were deposited in 48-well tissue culture plates (Corning® Costar®), in which the midguts were placed before the dome completely solidified. Matrigel™ provides structural support for cells because of its content of laminin, collagen IV, entactin, and heparin sulfate proteoglycan perlecan, among other factors (32), which are the main components of the mosquito midgut basal lamina secreted by epithelial cells (33). The support given by the hydrogel could be readily observed when embedding and culturing the midguts in Matrigel™. Microscopic observation of the *ex vivo* midguts cultured in medium showed some variation in the midgut organ integrity. Some midguts preserved their shape with a minimal amount of degradation over time, whereas others lost their integrity and progressively became misshapen, which has been noted before when working with mosquito tissue cultures (20). Of note, the hindgut was the region less prone to degradation. Such occurrence was improved greatly when using Matrigel™, as the midguts preserved their integrity better compared to the traditionally used culture medium. Therefore, the domes provided good support for the midgut organ, as they still presented hindgut peristalsis while keeping the tissue immobilized in the center of the well. However, infection of midguts kept in Matrigel domes by incubation with ZIKV was not successful. No viral RNA was detected in these midguts after infection (data not shown), as opposed to the midguts cultured traditionally (suspended in medium). Considering that the current route of infection diverges from a natural mosquito infection, the use of synthetic hydrogels could help to overcome such limitation while providing an adequate environment to culture the *ex vivo* mosquito midguts. For instance, the midguts embedded in a hydrogel are immobilized, which could allow the microinjection of virus directly into the midgut organ. Whether such approach could resemble the natural infection pattern of the midgut in mosquitoes warrants further investigation.

Another method tested consisted of using a foam substrate insert to line up the bottom of each well of the 96-well tissue culture plate filled with medium, as it was shown to be successful for the *ex vivo* culture of tick organs (14). Of note, the same foam substrate reported for tick organs could not be purchased due to its commercial unavailability. After dissection, mosquito midguts were laid into the sponge substrate and kept in culture. Unfortunately, the sponge substrate used had big pores that allowed the midguts to pass through the foam to the bottom of the well and posed difficulties in the washing steps; therefore, this method was also discarded. Of note, infection by incubation with ZIKV was successful in the midguts kept on foam substrate; however, traditionally culture midguts reached higher ZIKV RNA loads as measured by qRT-PCR (Supplementary material, Fig. S3). Furthermore, no organ controls, consisting of foam substrate (no midgut) that went along all infection steps, were implemented to measure any virus inoculum that could remain embedded in the substrate. These no organ controls were shown to retain a higher amount of the virus inoculum compared to the negative controls normally used (Paraformaldehyde [PFA] fixated midguts suspended in medium).

Therefore, the culture method that provided the best outcome in terms of handling, tissue preservation and permissiveness to arbovirus infection was to keep the mosquito midguts suspended in culture medium in a 96-well tissue culture plate with clear bottom (PerkinElmer©, USA). To reduce the risk of pipetting out the midguts during washing steps, a magnifying lamp was key to observe the midguts with the naked eye and avoid such problem. Lastly, carboxymethyl cellulose (CMC) 0.8% diluted in Leibovitz's L-15 medium was also employed and found to be suitable for long incubation periods (>3 days), as it provided a denser medium where the midguts seemed to preserve their shape better over time. However, viral RNA levels detected in the midguts incubated in CMC were significantly reduced at day 2 and 3 p.i. compared to traditionally *ex vivo* cultured midguts (Supplementary material, Fig. S4).
