## Supplementary Fig. S1 for "*Ex vivo* midgut cultures of *Aedes aegypti* are efficiently infected by mosquito-borne alpha- and flaviviruses"

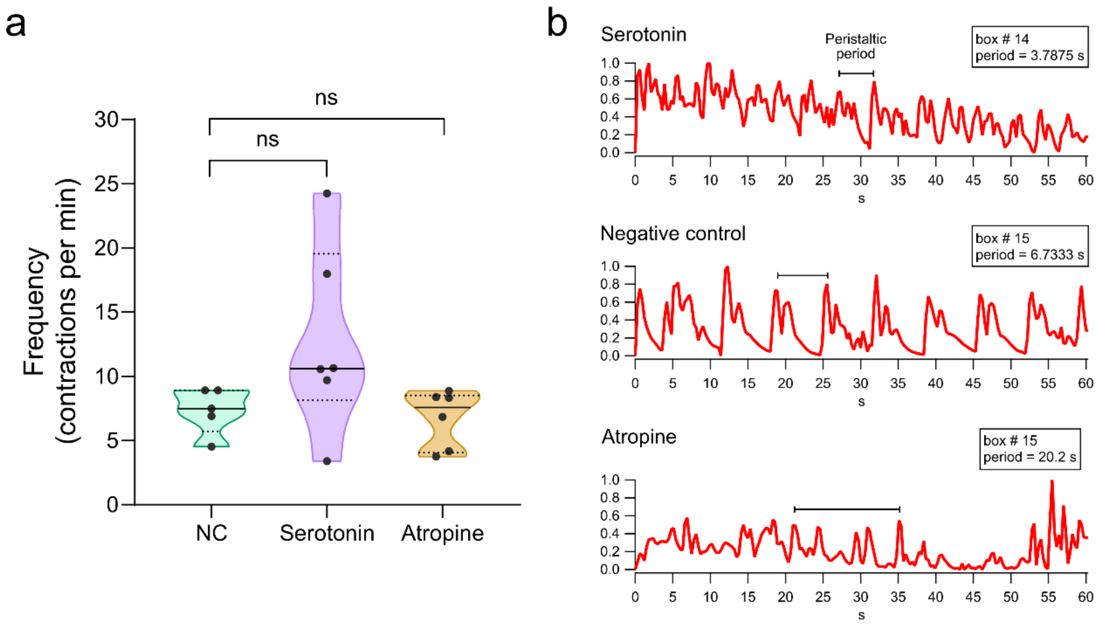


**Figure S1**. ***Ex vivo*** **cultured mosquito guts were not significantly affected by external incubation with serotonin or atropine.** **a**, Peristaltic period for each condition evaluated. NC stands for negative control, which represents the mock-exposed guts (untreated). Each point represents an individual mosquito midgut. The line represents the median value. The experiment was performed once. **b**, Representative output analysis for one midgut of each tested condition. The output corresponds to one region (box number) of the area of analysis (AOA) set for each gut. The peristaltic period in each output is representatively denoted as a black bar.
