## Supplementary Fig. S2 for "*Ex vivo* midgut cultures of *Aedes aegypti* are efficiently infected by mosquito-borne alpha- and flaviviruses"

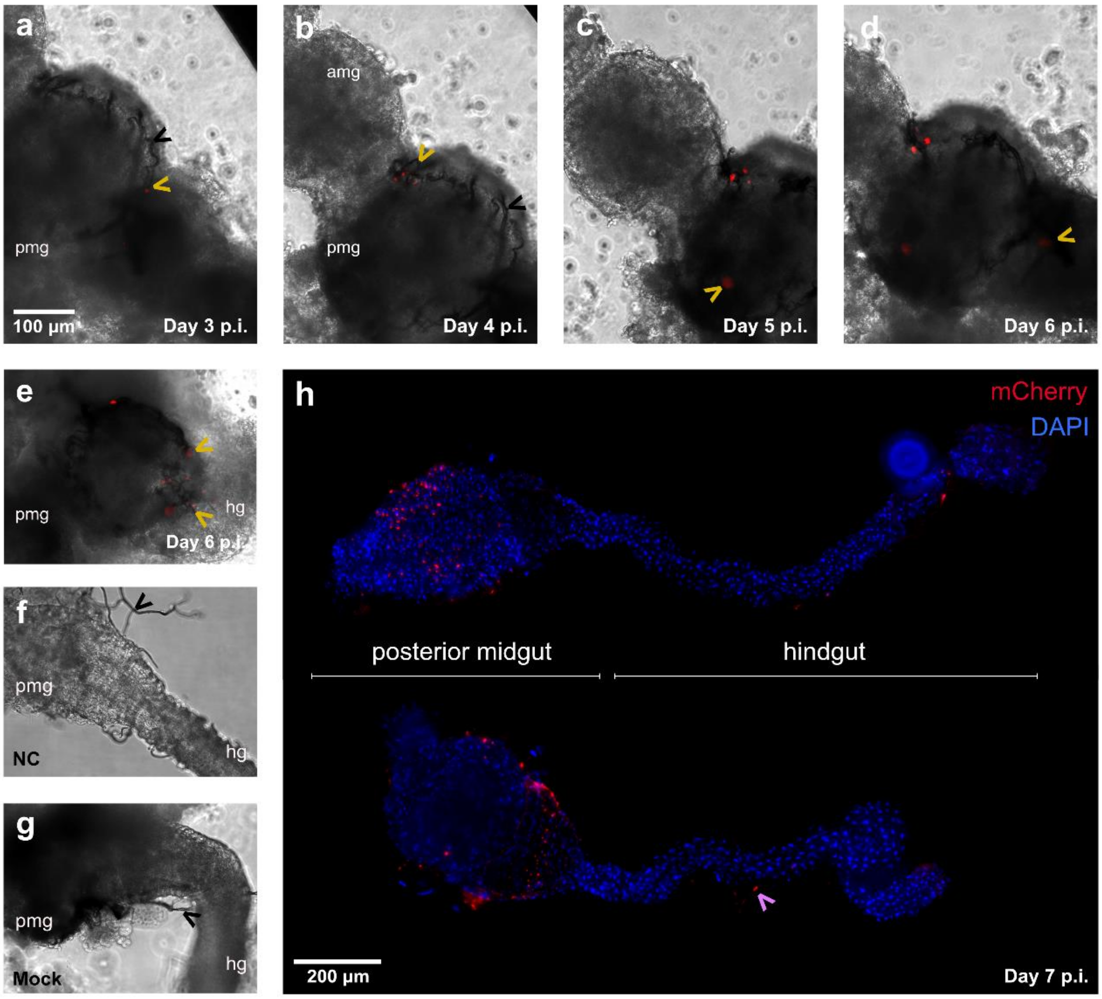


**Figure S2**. **Replication of DENV-2 expressing mCherry in the *ex vivo* cultured midguts. a**, **b**, **c**, **d**, Magnification, 20X. Overlay of bright field and red filter imaging. Panels display the DENV-2 infection and spread in the midguts as seen by the mCherry red signal on days 3 to 6 p.i. All four images correspond to the same organ imaged over the selected time points. The scale bar, represented by the white line, can be applied to panels a-g. **e**, mCherry expression in another biological replicate at day 6 p.i., showing several infection foci in the posterior midgut region. **f**, **g**, NC: negative control. No mCherry expression was detected in fixated or mock-infected midguts. **h**, mCherry signal in DENV-2 infected midgut, magnification, 10X. amg: anterior midgut, pmg: posterior midgut, hg: hindgut. Black arrows indicate the presence of tracheae. Yellow arrows indicate mCherry signal. Magenta arrow indicates mCherry signal present in tracheal tubes that remained in the hindgut.
