## Supplementary Fig. S3 for "*Ex vivo* midgut cultures of *Aedes aegypti* are efficiently infected by mosquito-borne alpha- and flaviviruses"

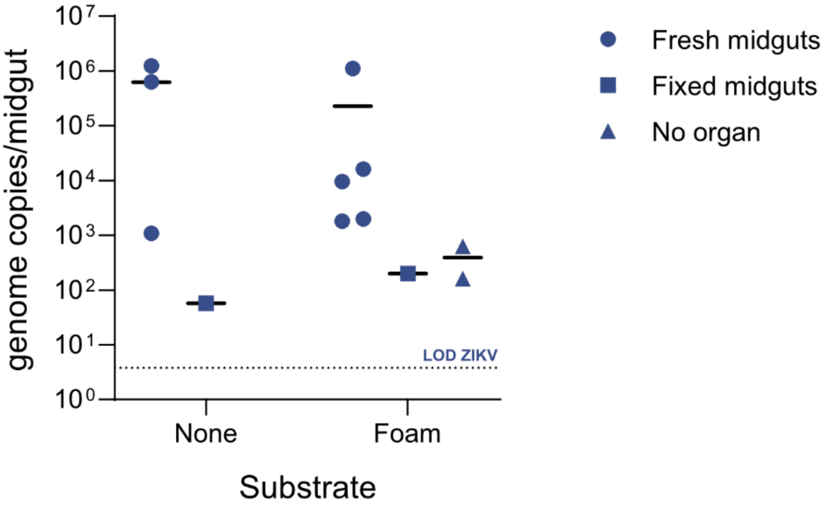


**Figure S3**. **ZIKV RNA loads in the mosquito midguts when cultured in medium or when using a foam substrate for support.** RNA levels were quantified at day 7 p.i. by qRT-PCR. The black line represents the mean value. Fresh midguts: alive midguts, negative control: PFA fixated midguts, no organ: foam substrate only. LOD: limit of detection of the assay.
