## Supplementary Fig. S4 for "*Ex vivo* midgut cultures of *Aedes aegypti* are efficiently infected by mosquito-borne alpha- and flaviviruses"

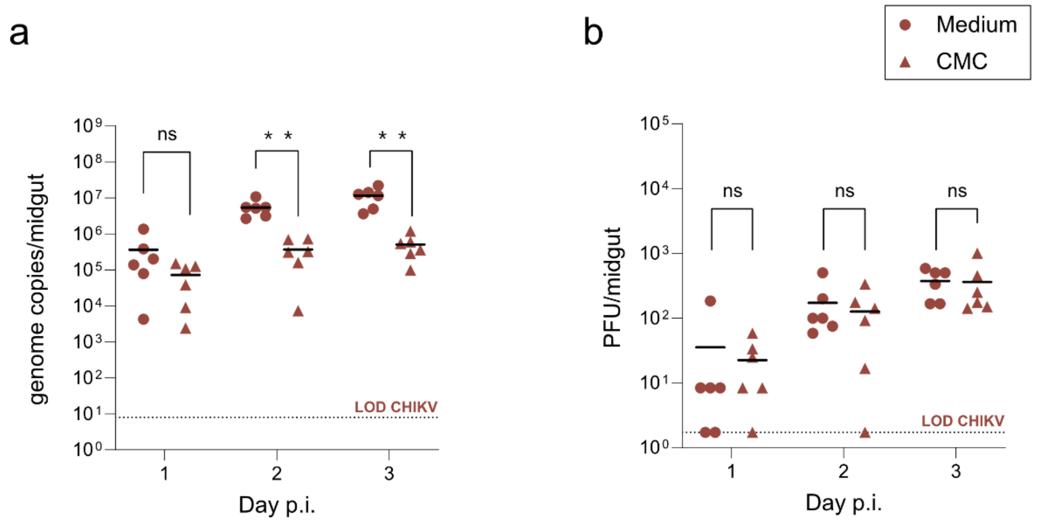


**Figure S4**. **CHIKV replication kinetics in mosquito midguts cultured in medium and in carboxymethylcellulose (CMC). a**, Viral RNA levels were quantified at day 1, 2 and 3 p.i. by means of qRT-PCR. **b**, Infectious virus loads in the mosquito midguts were quantified by means of plaque assay. Each dot represents an individual midgut organ. The black line represents the mean value. Statistical significance was assessed with a Mann-Whitney test. Significantly different values are indicated by asterisks: **, P<0.005. ns: not significant. LOD: Limit of detection of the corresponding assay.
